# BLAST: A blue light-assisted secretion toolkit tunable by reversible protein-protein interactions

**DOI:** 10.64898/2026.03.30.715452

**Authors:** Meiying Shen, Seunghwan Lee, Kyuye Song, Mingguang Cui, Dongmin Lee

## Abstract

Precise control over protein secretion is essential for programming intercellular communication and coordinating complex physiological responses. However, conventional methods relying on transcriptional regulation or chemical induction often lack the spatiotemporal precision and reversibility required to mimic endogenous signaling dynamics. Here, we present the Blue Light-Assisted Secretion Toolkit (BLAST), a genetically encoded system that orchestrates protein release from the endoplasmic reticulum via light-tunable protein-protein interactions. BLAST comprises two complementary modules utilizing both light-induced iLID/SspB association (a-BLAST) and LOV2/Zdk1 dissociation (d-BLAST). Both modules harness the highly conserved RXR motif to enforce strict ER confinement in the dark state. Most importantly, by utilizing non-destructive steric masking rather than enzymatic cleavage, BLAST achieves unprecedented temporal resolution with strict reversibility. We demonstrate that both systems can be repeatedly toggled ON and OFF, instantaneously arresting cargo release upon light withdrawal to generate highly controlled, pulsatile secretion profiles. Leveraging this dynamic control, we successfully achieved the rapid, robust, and light-triggered secretion of complex therapeutic proteins, including insulin and interleukin-12. By bypassing transcriptional delays and irreversible activation steps, BLAST provides a generalized, plug-and-play platform for the on-demand delivery of therapeutic proteins, significantly expanding the optogenetic toolbox for synthetic biology and cell-based therapies.

## 1. Introduction

Secreted proteins serve as the primary currency of intercellular communication, orchestrating diverse physiological processes ranging from immune responses to synaptic transmission^[1–3]^. The dysregulation of these secretory pathways is often linked to severe pathological conditions^[4–6]^, underscoring the need for precise therapeutic interventions. However, natural secretion is a highly dynamic event, often occurring in pulsatile or localized bursts^[7–9]^. Recapitulating these complex profiles for therapeutic and synthetic biology applications requires tools that allow for the precise spatiotemporal control of protein release, surpassing the capabilities of constitutive secretion.

Current strategies for controlling protein secretion predominantly rely on transcriptional regulation^[10–17]^. While effective for long-term expression, these methods are inherently limited by the slow kinetics of transcription and translation, requiring at least 4–6 h to reach therapeutic levels^[18]^. To achieve faster kinetics, post-translational retention systems have been developed. Early approaches, such as the chemically inducible RUSH (retention using selective hooks)^[19]^ or protease-dependent strategies like RELEASE (Retained Endoplasmic Cleavable Secretion)^[20]^, membER/lumER^[21]^, and POSH (protease-mediated post-translational switch)^[22]^ significantly improved response times compared to transcriptional control. However, these methods often depend on exogenous ligands that are difficult to wash out and lack the dynamic reversibility required for strict ON/OFF control.

Subsequently, the development of optogenetic tools, such as optoPOSH^[22]^ and optoPASS^[23]^, introduced a new level of spatiotemporal resolution. By employing pMag and nMag dimerization modules to reconstitute a split protease upon illumination^[24,25]^, optoPOSH and optoPASS successfully triggers protein release. However, because this system fundamentally relies on irreversible proteolytic cleavage, it functions as a single-activation switch, preventing the reversible toggling of secretion required to mimic dynamic physiological signals.

To overcome these limitations, we developed the Blue Light-Assisted Secretion Toolkit (BLAST), a genetically encoded platform that exerts rapid and reversible control over protein secretion. Distinct from previous cleavage-based methods, BLAST strategically harnesses the RXR (Arginine-X-Arginine) motif—a stringent sorting signal utilized by native channels—to enforce strict spatial confinement within the ER^[26,27]^. By coupling this robust RXR-mediated retention with light-tunable protein-protein interactions (PPIs), BLAST employs non-destructive steric masking rather than enzymatic cleavage. This mechanism not only bypasses the central dogma lag to enable the immediate release of pre-synthesized cargo, but crucially allows for the instantaneous cessation of secretion upon light withdrawal. We engineered two complementary systems: an associative module (a-BLAST) based on Light-Inducible Dimer (iLID)/stringent starvation protein B (SspB) dimerization^[28,29]^ and a dissociative module (d-BLAST) based on Light-Oxygen-Voltage sensing domain 2 (LOV2)/Zdk1 dissociation^[30]^. Here, we demonstrate that BLAST provides a robust, plug-and-play solution for the spatiotemporally precise delivery of various cargo proteins, offering a generalized method to program intercellular signaling with blue light.

## 2 Results

### 2.1 Design and characterization of the associative BLAST system

To engineer a precision tool for light-regulated protein secretion, we designed the associative Blue Light-Assisted Secretion Toolkit (a-BLAST) (Fig. 1a). This system exploits the 470 nm blue light-dependent heterodimerization between the iLID and its binding partner, SspB^[28,29]^. The a-BLAST system comprises two modular components: an ER-anchored cargo module and a cytosolic regulator module. The prototype cargo module is designed with an N-terminal protein of interest (POI), followed by a Furin cleavage site, a transmembrane domain (TM), an SspB sequence, and a C-terminal RXR ER-retention motif^[27]^. In the dark state, the cargo is strictly confined within the ER due to the exposed RXR motif, which is recognized by the COPI retrieval machinery^[31]^. Upon blue light illumination, the cytosolic iLID regulator binds to the SspB moiety on the cargo. We hypothesized that this light-induced association would sterically mask the RXR motif^[27,32]^, non-destructively overriding the retention signal and permitting the cargo’s exit to the trans-Golgi network. Subsequently, endogenous Furin proteases within the Golgi cleave the processing site, resulting in the secretion of the mature POI.

**Figure 1.**
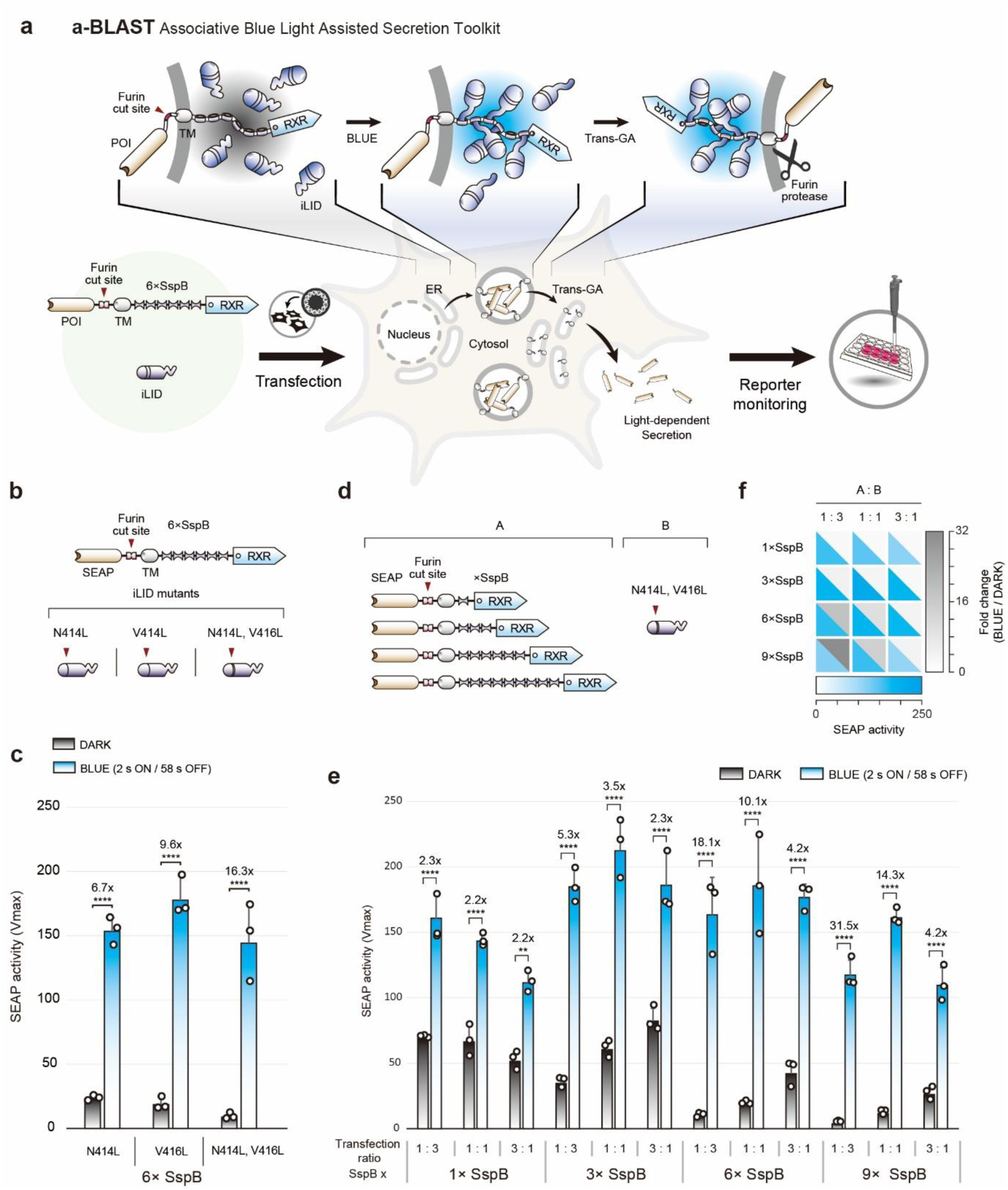
Development of the associative Blue Light-Assisted Secretion Toolkit (a-BLAST). (a) Schematic illustration of the a-BLAST mechanism. The a-BLAST system consists of two parts: a cargo module and an associative regulator module. The cargo module includes the POI, a Furin cut site, a TM domain, SspB repeats, and an RXR-type ER retention sequence. The associative regulator comprises iLID, which binds to SspB under blue light. Exposure to blue light triggers SspB-iLID dimerization, which sterically interferes with the RXR-mediated ER retention. Consequently, the cargo is transported to the trans-Golgi network, where it is cleaved by endogenous Furin protease to release the POI. (b) Construct configurations of a-BLAST components: the cargo (upper) and the associative regulator (lower). The cargo features SEAP as a reporter, a TM domain, a Furin cleavage site, 6× SspB, and an RXR motif. The regulator variants include iLID mutants with delayed recovery kinetics. (c) Comparative analysis of blue light-dependent protein secretion based on kinetic modifications of iLID. The iLID double mutant (N414L, V416L) exhibited the highest fold change (16.3-fold). Blue light (470 nm) was applied for 24 h with a duty cycle (2 s ON / 58 s OFF). (d) Plasmid library demonstrating sequential incrementation of SspB repeats (A: 1×, 3×, 6×, and 9×) and the dual-mutant iLID (B). (e) Histogram of SEAP activity and fold changes based on varying transfection ratios and SspB valency. The configuration using 6× SspB and a 1:3 transfection ratio yielded the most robust fold change (18.1-fold). (f) Heatmap summarizing blue light-induced SEAP activity (white-to-blue gradient) and fold changes (white-to-gray gradient). Open circles represent individual measurements from three biologically independent samples. Data are presented as means ± S.D. Significance was assessed using one-way ANOVA followed by Tukey’s multiple comparisons test. (ns = not significant, \**P* < 0.05, \*\**P* < 0.01, \*\*\**P* < 0.001, \*\*\*\**P* < 0.0001).

To optimize the dynamic range of this prototype, we utilized Secreted Embryonic Alkaline Phosphatase (SEAP) as a reporter system. We compared the wild-type iLID with three mutants (N414L, V416L, and N414L/V416L) known to exhibit distinct binding affinities (Fig.1b, Supplementary Fig. 1a)^[29]^. Consistent with previous characterizations of iLID variants^[29]^, the double mutant (N414L/V416L) exhibited the highest dynamic range and was selected for all subsequent optimizations (Fig.1c, Supplementary Fig. 1c).

Next, to maximize the signal-to-noise ratio, we focused on optimizing the C-terminal topology of the retention module. Since RXR-mediated retention is sensitive to its spatial distance from the membrane and the surrounding steric context^[31]^, we hypothesized that altering the spacer length via SspB concatemerization would critically influence retention efficiency. We generated a library of cargo variants containing 1× to 9× tandem repeats of SspB (Fig. 1d). Through a comprehensive screen of SspB valencies and plasmid transfection ratios (Cargo: Regulator), we evaluated the secretory performance of each configuration (Fig. 1e, f). Although the 9× SspB variant at a 1:3 ratio exhibited the highest apparent fold-change (31.5-fold), its absolute maximum secretion level under blue light was relatively diminished. This reduction is likely attributable to the structural burden and reduced translation or folding efficiency imposed by the excessively large tandem repeats. Therefore, we selected the 6× SspB variant coupled with a 1:3 transfection ratio as the optimal configuration. This setup provided the ideal balance between stringent dark-state retention and a robust, high-capacity protein output, achieving a 16.3-fold increase in SEAP secretion (Fig. 1c). These results demonstrate that precisely tuning both the steric environment of the retention motif and the stoichiometry of the system components is critical for achieving accurate secretory control.

### 2.2 Benchmarking the reversible retention mechanism against proteolytic release

To rigorously benchmark the efficiency of the reversible steric masking strategy of a-BLAST against established irreversible proteolytic cleavage systems, we compared BLAST with the RELEASE system^[20]^. RELEASE utilizes a split protease to cleave an ER-retention signal (KKMP), a mechanism that typically requires close proximity to the ER membrane for efficient proteolytic processing^[31]^. To ensure a direct, fair mechanistic comparison independent of the stimulus modality (light versus drug), we engineered a chemogenetic analog of a-BLAST (chemo-BLAST) by substituting the iLID/SspB optogenetic pair with the rapamycin-inducible FRB/FKBP dimerization system (Supplementary Fig. 2a)^[33]^.

We first optimized the chemo-BLAST architecture by varying the valency of the FRB-fused retention module (1×, 2×, 4×, and 6×). In contrast to the optogenetic a-BLAST (which preferred a 6× configuration), the chemo-BLAST variant exhibited the highest rapamycin-dependent dynamic range with 4× FRB repeats (Supplementary Fig. 2b). Subsequent optimization of the transfection ratio identified that a 1:3 (Cargo: Regulator) ratio yielded a maximal 11.3-fold induction (Supplementary Fig. 2c). Next, we performed a head-to-head comparison between the optimized chemo-BLAST and the RELEASE system (employing both TEVp and TVMVp split proteases). The results unequivocally demonstrated that chemo-BLAST significantly outperformed the RELEASE system in terms of overall secretion efficiency (Supplementary Fig. 2d). Notably, chemo-BLAST exhibited markedly superior kinetics, inducing significant protein secretion within just 4 h of rapamycin treatment, whereas the RELEASE system showed a pronounced temporal delay. These findings strongly suggest that the non-destructive steric masking mechanism of BLAST allows for immediate cargo exit upon inducer binding, effectively bypassing the kinetic bottleneck associated with the proteolytic cleavage step required by systems like RELEASE.

### 2.3 Design and characterization of the dissociative BLAST system

To provide a complementary mode of regulation characterized by rapid response kinetics, we developed the dissociative BLAST (d-BLAST) system, inspired by the Light-Oxygen-Voltage sensing domain 2 Trap and Release of Protein (LOVTRAP) optogenetic tool (Fig. 2a)^[30]^. Unlike the associative a-BLAST, this system operates via light-induced dissociation. The d-BLAST architecture consists of an ER-anchored cargo module (POI-Furin-TM-mCherry-LOV2) and a cytosolic regulator module (Zdk1-RXR) (Fig. 2a). In the dark, the Zdk1 domain of the regulator binds tightly to the LOV2 domain on the cargo. This interaction transiently recruits the RXR motif to the cargo, effectively hijacking the COPI retrieval machinery to strictly retain the protein within the ER. Upon blue light irradiation, the Jα helix of LOV2 unfolds, causing the rapid dissociation of the Zdk1-RXR complex. This instantaneous removal of the retention signal permits the cargo to escape the ER and proceed through the secretory pathway.

**Figure 2.**
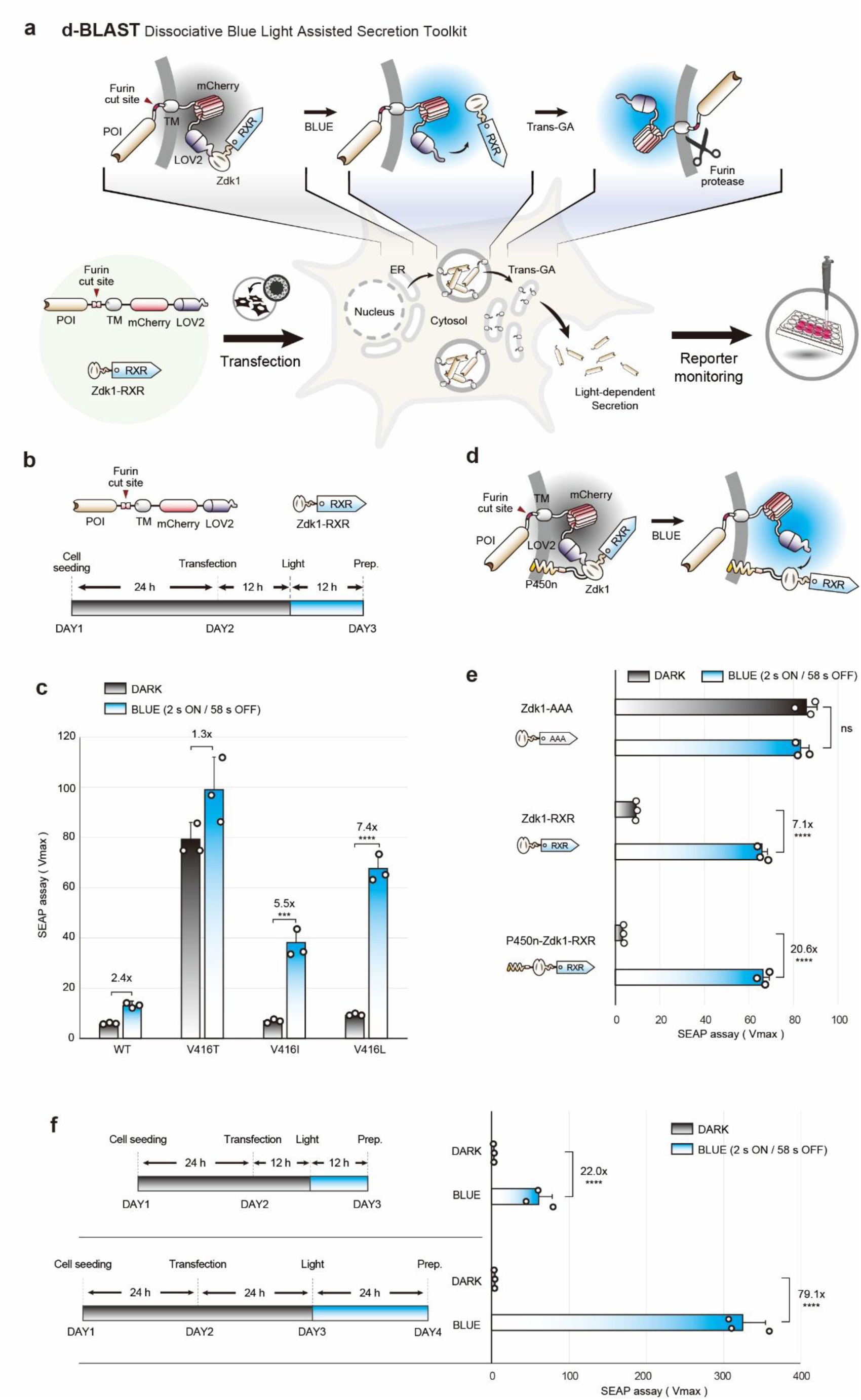
Development of the dissociative-BLAST (d-BLAST) system. (a) Schematic diagram of the d-BLAST system. d-BLAST consists of two components: a cargo module and a dissociative regulator module. The cargo module contains a protein of interest (POI; e.g., SEAP), a Furin cleavage site, a transmembrane (TM) domain, an mCherry spacer, and a LOV2 domain. The dissociative regulator consists of Zdk1 fused to a C-terminal RXR ER retention motif. In the dark, Zdk1 binds tightly to LOV2, recruiting the RXR motif to the cargo and retaining it in the endoplasmic reticulum (ER). Blue light induces the dissociation of Zdk1-RXR from the cargo, allowing the POI to be transported to the trans-Golgi network (Trans-GA), where it is cleaved by native Furin protease and secreted. (b) Structural maps and experimental timeline. Schematic representation of the cargo and dissociative regulator constructs, along with a 3-day experimental schedule. (c) Kinetic optimization using LOV2 mutants. Comparison of different LOV2 domain mutants (WT, V416T, V416I, and V416L) to control protein secretion. The V416L mutant, which has the slowest reversion kinetics, exhibited the highest fold change in SEAP secretion. (d) Schematic of ER membrane-localized regulator. To reduce basal leakage, the dissociative regulator was anchored to the ER membrane by fusing a Cytochrome P450 signal-anchor sequence (P450n) to the N-terminus of Zdk1-RXR. (e) Influence of regulator localization on secretion. Comparison of SEAP secretion levels using different Zdk1 variants. Anchoring the regulator (P450n-Zdk1-RXR) to the ER membrane significantly reduced dark-state leakage compared to the cytosolic version, improving the signal-to-noise ratio to 20.6-fold. The Zdk1-AAA mutant lacks retention function, resulting in high, light-independent secretion. (f) Optimization of experimental schedules. Comparison of d-BLAST performance between a 3-day and 4-day schedule. The 4-day schedule significantly increased the total level of light-induced protein secretion (79.1-fold). Open circles represent individual measurements from three biologically independent samples. Data are presented as means ± S.D. Significance was assessed using one-way ANOVA followed by Tukey’s multiple comparisons test (ns = not significant, \**P* < 0.05, \*\**P* < 0.01, \*\*\**P* < 0.001, \*\*\*\**P* < 0.0001).

Crucially, because the RXR-mediated retention mechanism requires a specific spatial distance from the membrane to function effectively, we incorporated mCherry as a spacer to optimally position the RXR motif away from the ER membrane^[31]^. Validating this design, constructs lacking the mCherry spacer exhibited severe dark-state leakage (Supplementary Fig. 3), confirming the spacer’s structural necessity for strict spatial retention. Having established the optimal cargo architecture containing the mCherry spacer, we next optimized the photocycle kinetics to enable efficient secretion while minimizing phototoxicity. We screened LOV2 mutants (V416T, V416I, V416L) with varying dark reversion rates, hypothesizing that slow-reverting variants would maintain the dissociated (ON) state longer under pulsed illumination. Using a 12 h stimulation protocol (2 s ON/58 s OFF) (Fig. 2b), we confirmed that the V416L mutant, which possesses the slowest reversion kinetics, yielded the highest fold-change in SEAP secretion (Fig. 2c).

However, a persistent challenge with dissociative systems is basal leakage in the dark state due to the incomplete capture of the cargo by cytosolic regulators. To address this, we engineered a membrane-tethered regulator by fusing the Zdk1-RXR module to the N-terminal signal anchor of cytochrome P450 (P450n)^[34,35]^ (Fig. 2d). We reasoned that immobilizing the regulator on the ER membrane would drastically increase its local effective concentration around the cargo, thereby enforcing stricter retention in the dark. Consistent with our design rationale, the P450-anchored regulator substantially minimized dark-state basal leakage compared to the cytosolic version, boosting the optogenetic dynamic range from 7.1-fold to an impressive 20.6-fold (Fig. 2e). To further maximize protein yield, we optimized the production timeline. Extending the protocol from a 3-day schedule (12 h expression/12 h light) to a 4-day schedule (24 h expression/24 h light) resulted in a remarkable 79.1-fold increase in cumulative protein secretion (Fig. 2f).

### 2.4 Kinetic profiling and benchmarking of BLAST against proteolytic switches

To precisely define the temporal resolution of our toolkit, we profiled the secretion kinetics of both a-BLAST and d-BLAST. HEK293T cells expressing optimized constructs were subjected to blue light or dark conditions for varying durations (0 to 24 h) starting 24 h post-transfection (Fig. 3a, b). Strikingly, d-BLAST induced statistically significant SEAP secretion within just 1 h (3.5-fold), whereas a-BLAST required 4 h of illumination to achieve significant release (5.2-fold) (Fig. 3c). Comparing the two, d-BLAST exhibited superior rapid-response kinetics, achieving a 2.2-fold increase in secretion after only 30 m of illumination. However, this high sensitivity was accompanied by a time-dependent increase in basal secretion in the dark, likely attributable to the hypersensitivity of the dissociative mechanism to ambient light exposure during sample handling. In contrast, a-BLAST maintained exceptionally low background levels because the RXR retention motif is inherently incorporated into the cargo module itself, resulting in a robust signal-to-noise ratio over prolonged durations. Thus, the toolkit offers a strategic trade-off: d-BLAST for applications requiring immediate release, and a-BLAST for those demanding minimal background.

**Figure 3.**
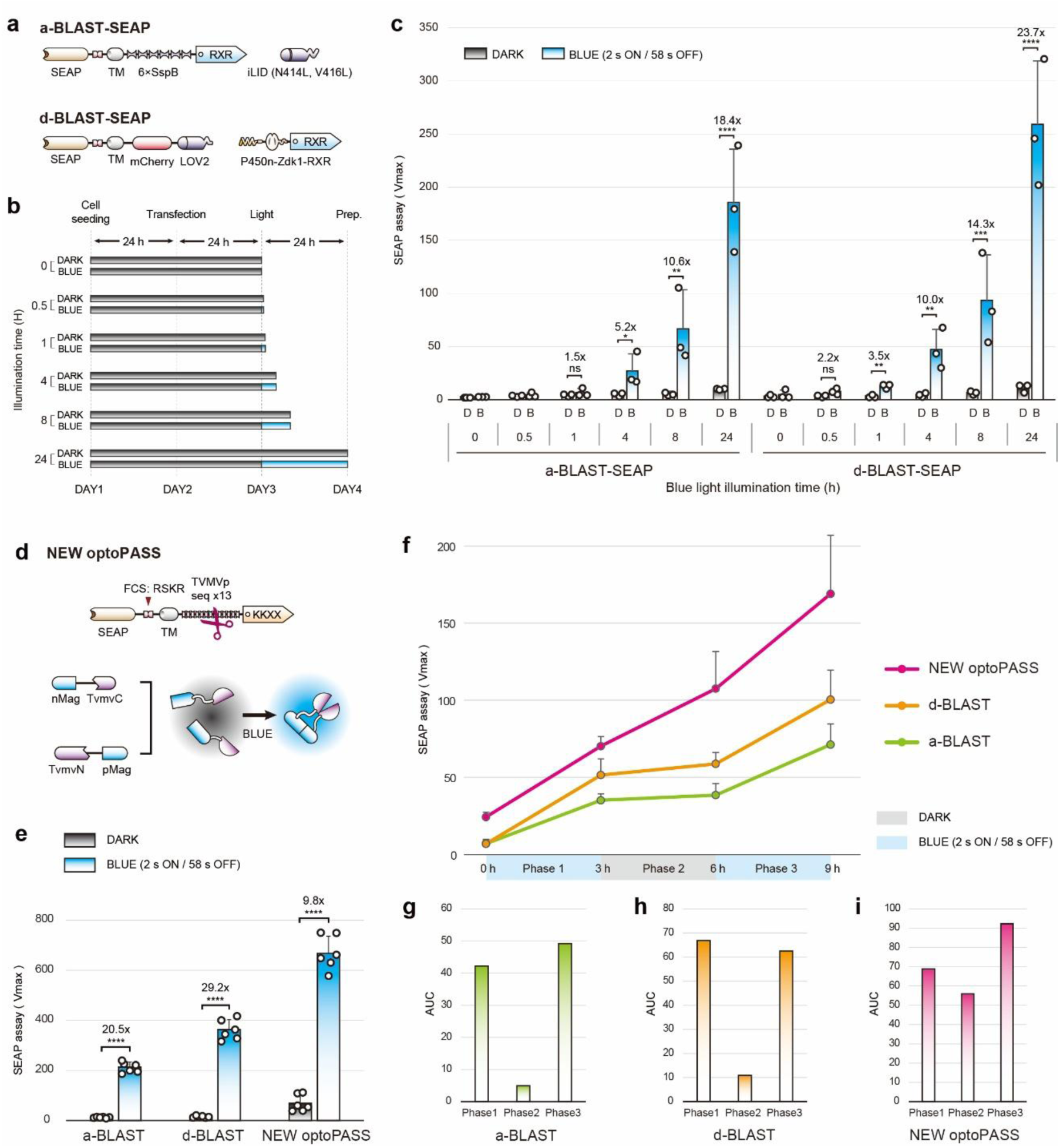
Characterization of a-BLAST and d-BLAST depending on blue light duration and reversibility. (a) Structural diagrams for kinetic characterization. Schematic representation of optimized SEAP-a-BLAST (upper) and SEAP-d-BLAST (lower). The a-BLAST system utilizes 6× SspB and the iLID (N414L, V416L) double mutant. The d-BLAST system employs the LOV2 domain and the P450n-anchored Zdk1-RXR regulator. (b) Experimental timeline. Cells were subjected to varying intervals of blue light illumination (0, 0.5, 1, 4, 8, and 24 h) starting 24 h post-transfection. (c) Summary graphs showing SEAP secretion levels for a-BLAST (left) and d-BLAST (right) across different illumination times. Both systems exhibit light-dependent secretion, with d-BLAST showing rapid release within 1 h (3.5-fold) and a 2.2-fold increase even at 0.5 h. a-BLAST shows robust, significant secretion starting at 4 h (5.2-fold). (d) Schematic of the NEW optoPASS benchmarking construct. The original Furin cleavage site and TM domain were replaced with those used in BLAST. Blue light triggers pMag-nMag dimerization, reconstituting split protease (TVMVp) activity to excise the KKMP retention motif. (e) Comparison of SEAP secretion and signal-to-noise ratios between a-BLAST, d-BLAST, and NEW optoPASS. While NEW optoPASS shows higher absolute secretion, the BLAST systems demonstrate superior dynamic ranges (20.5-fold for a-BLAST, 29.2-fold for d-BLAST) due to lower dark-state leakage compared to NEW optoPASS (9.8-fold). (f) Cumulative SEAP secretion profiles demonstrating the reversibility of the systems over a 9-h alternating light/dark cycle (3 h ON / 3 h OFF / 3 h ON). Both a-BLAST (green line) and d-BLAST (orange line) generate strictly controlled staircase profiles, instantaneously halting secretion during the dark phase. In contrast, NEW optoPASS (magenta line) exhibits continuous, irreversible leakage. (g–i) Area under the curve (AUC) quantification of secretion levels during each 3-h phase for (g) a-BLAST (green bars), (h) d-BLAST (orange bars), and (i) NEW optoPASS (magenta bars). The strictly minimal AUC during Phase 2 (Dark) for both BLAST systems definitively confirms robust ON/OFF control compared to the cleavage-based system. Open circles represent individual measurements from three biologically independent samples. Data are presented as means ± S.D. Statistical significance was assessed using one-way ANOVA followed by Tukey’s multiple comparisons test (ns = not significant, \**P* < 0.05, \*\**P* < 0.01, \*\*\**P* < 0.001, \*\*\*\**P* < 0.0001).

We further validated the universality of BLAST across diverse mammalian cell lines. Both systems functioned robustly in HeLa, BHK-21, and HepG2 cells (Supplementary Fig. 4a-d), confirming that BLAST relies on the conserved secretory machinery of mammalian cells and is applicable to a wide range of biological contexts.

Next, we benchmarked BLAST against optoPASS^[23]^, a recently developed state-of-the-art system that triggers secretion via the light-induced reassembly of a split protease. While optoPASS also utilizes blue light, it relies on an influenza virus-derived Furin cleavage site (FCS: RRRKKR/GL), whereas our BLAST employs a Sindbis virus-derived sequence (SGRSKR/SV). To ensure a rigorous, variable-controlled comparison, we engineered a chimeric variant named NEW optoPASS, in which the FCS and transmembrane (TM) domains were replaced with those from BLAST (Fig. 3d). Under identical conditions, although NEW optoPASS yielded higher absolute secretion levels, it suffered from significant basal leakage in the dark, likely due to the irreversible nature of proteolytic escape. Conversely, BLAST demonstrated a significantly superior signal-to-noise ratio (Fig. 3e). This conclusion was further supported by reciprocal domain-swapping experiments (Supplementary Fig. 5).

To rigorously demonstrate the dynamic, multi-toggling capability of our toolkit—a critical requirement for mimicking pulsatile physiological signaling—we evaluated the reversibility of BLAST in direct comparison to NEW optoPASS. We hypothesized that the non-destructive, reversible masking of the ER retention signal in BLAST would enable the immediate cessation of cargo exit upon light withdrawal, offering superior ON/OFF control. To test this, we tracked cumulative SEAP secretion over a 9-hour period under an alternating light/dark cycle (3 h ON / 3 h OFF / 3 h ON). Supernatants were sampled at the corresponding time points (0, 3, 6, and 9 h). This schedule allowed us to precisely monitor secretion profiles during illumination windows, while evaluating dark-state retention by comparing cumulative levels before and after the dark period.

Strikingly, the resulting profiles revealed a profound mechanistic divergence, highlighting the fundamental difference between irreversible cleavage and reversible masking (Figure 3f). The NEW optoPASS system exhibited a continuous, unchecked increase in cumulative secretion; a substantial accumulation of SEAP was detected at 6 h compared to the 3 h time point (Phase 2), demonstrating that proteolytic escape completely fails to terminate protein release during the dark phase. In stark contrast, both a-BLAST and d-BLAST generated highly controlled “staircase” secretion profiles. While blue light illumination robustly triggered secretion, cumulative SEAP levels firmly plateaued during the intervening dark phase (between 3 h and 6 h), confirming that the re-establishment of steric masking instantaneously arrests further cargo release. This strict ON/OFF control is further corroborated by the area under the curve (AUC) analysis (Figure 3g–i), which clearly quantifies the minimal secretion during the dark phase for both BLAST systems compared to the continuous leakage of NEW optoPASS. Collectively, these results confirm that unlike irreversible “one-off” protease-dependent switches, the reversible retention mechanism of BLAST permits strict, repeatable multi-toggling, providing the unprecedented temporal resolution required for advanced synthetic biology applications.

### 2.5 Visualization of intercellular signaling and high-precision spatial control

To visualize the intercellular transfer of secreted proteins, we engineered a synthetic paracrine signaling circuit utilizing EGFP as a fluorescent cargo. We established a sender-receiver co-culture system: sender cells were transfected with the BLAST-EGFP module, while receiver cells expressed a membrane-tethered GFP-nanobody designed to capture secreted EGFP on their surface (Fig. 4a)^[36]^. Sender and receiver cells were seeded on separate coverslips and then co-cultured in close proximity, with the receiver coverslip inverted over the sender monolayer (Fig. 4b). Following blue light illumination, we analyzed the intercellular transfer via confocal microscopy. In the dark conditions, receiver cells showed no observable green fluorescence, confirming the tight retention of EGFP within the sender cells. In contrast, under blue light stimulation, distinct EGFP signals were detected on the plasma membrane of receiver cells, strongly co-localizing with the mCherry surface marker (Fig. 4c). This result visually confirms that BLAST facilitates the efficient secretion and subsequent intercellular transfer of functional cargo proteins, which can be specifically recognized by neighboring target cells.

**Figure 4.**
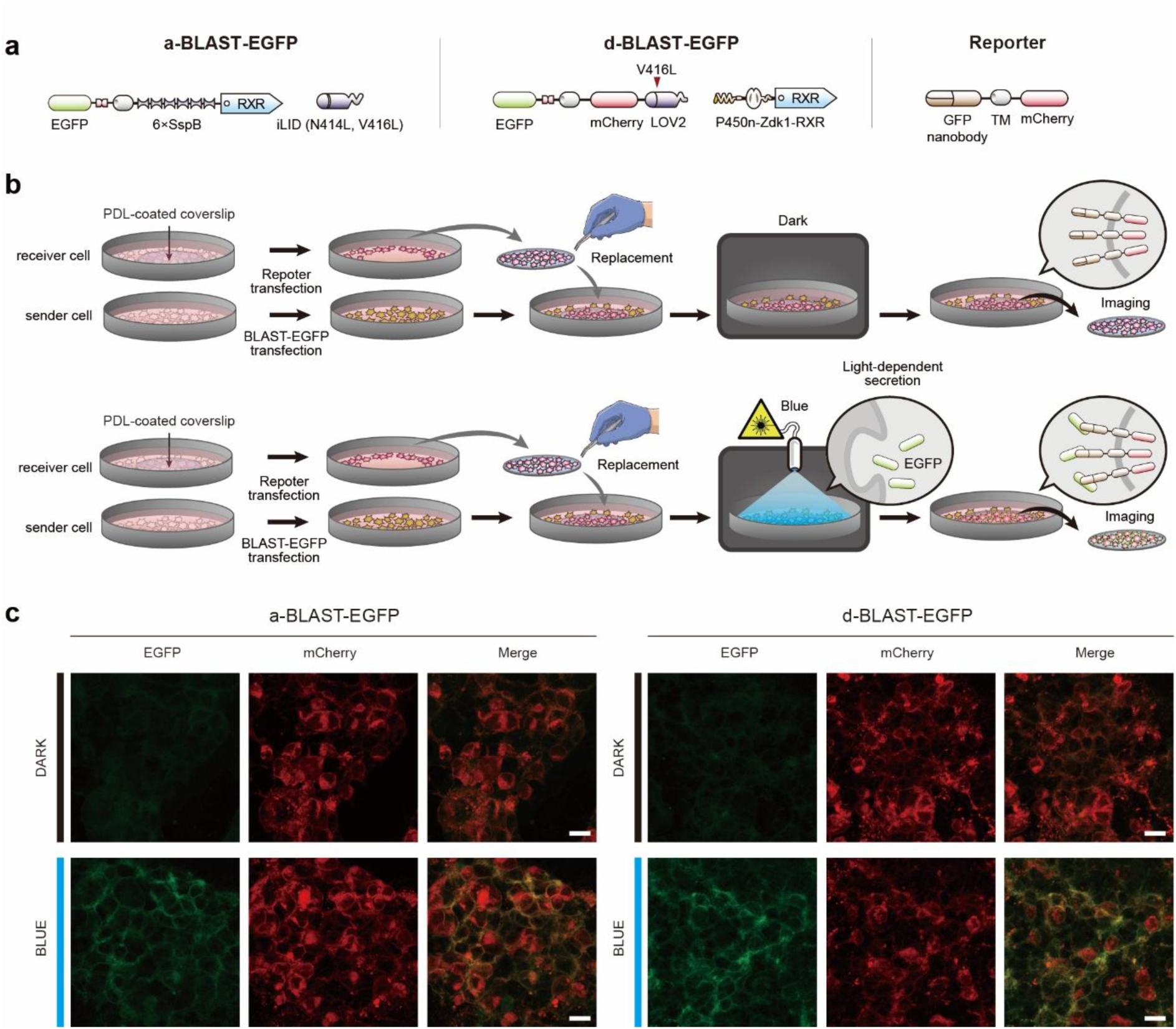
Visualization of a-BLAST and d-BLAST with EGFP and GFP nanobody. (a) Plasmid configurations of the BLAST-EGFP systems and the reporter. Schematic of EGFP-fused a-BLAST (left) and d-BLAST (middle). For visualization, the SEAP reporter was replaced with EGFP. The receiver reporter construct (right) consists of a membrane-tethered GFP-nanobody (GBP) fused to an mCherry transfection marker. (b) Schematic illustrating the sequential co-culture procedure. Sender cells (transfected with BLAST-EGFP) and receiver cells (transfected with the GBP reporter) were prepared on separate coverslips. The receiver coverslip was then transferred onto the sender cell plate. For the light-induced group, blue light stimulation triggers EGFP secretion and subsequent capture by the nanobody on the receiver cells, while dark conditions strictly retain EGFP within the sender cells. (c) Representative confocal images of a-BLAST and d-BLAST signaling. Confocal microscopy images showing receiver cells in dark (upper) and blue light (lower) conditions. The green signal (EGFP) indicates the successful capture of secreted EGFP by the nanobody on the plasma membrane of receiver cells. The red signal (mCherry) confirms the presence and localization of the reporter module on the cell surface. Scale bar: 20 μm.

Furthermore, we demonstrated the capacity of BLAST for spatially precise control using a photomasking strategy. Monolayers of HEK293T cells expressing a-BLAST-EGFP or d-BLAST-EGFP were illuminated through a 2-mm slit mask (Supplementary Fig. 6a, b). To enable rigorous quantitative analysis, the d-BLAST system utilized its intrinsic mCherry spacer as a reference marker. In contrast, because the a-BLAST cargo lacks this structural component, we co-transfected mCherry to serve as a normalization control. As anticipated, we observed a spatially defined negative photopatterning effect. Specifically, the targeted region exposed to blue light exhibited a pronounced reduction in intracellular EGFP fluorescence, while the signal from the mCherry reference marker remained completely unchanged compared to the surrounding dark areas (Supplementary Fig. 6c, d). This localized depletion of the EGFP cargo, coupled with the stable mCherry signal, clearly indicates that protein secretion was actively triggered only within the illuminated zone without affecting overall cell viability or basal expression, thereby confirming that BLAST enables high-resolution spatiotemporal control of protein release.

### 2.6 Light-triggered release of therapeutic proteins: Insulin and Interleukin-12

To evaluate the clinical potential and translatability of BLAST, we tested its ability to regulate the secretion of two medically significant proteins: human preproinsulin and the immunomodulatory cytokine interleukin-12 (IL-12). First, we engineered a preproinsulin cargo module by flanking the C-peptide with two Furin cleavage sites to ensure its maturation into active insulin via the endogenous secretory pathway (Fig. 5a)^[37]^. To facilitate efficient processing, we co-expressed Furin protease in HEK293T cells and quantified insulin release by measuring secreted C-peptide levels via enzyme-linked immunosorbent assay (ELISA). Consistent with our SEAP reporter results, both a-BLAST and d-BLAST induced significant insulin secretion within 2 h of illumination, exhibiting a clear dependency on the duration of light exposure (Fig. 5b).

**Figure 5.**
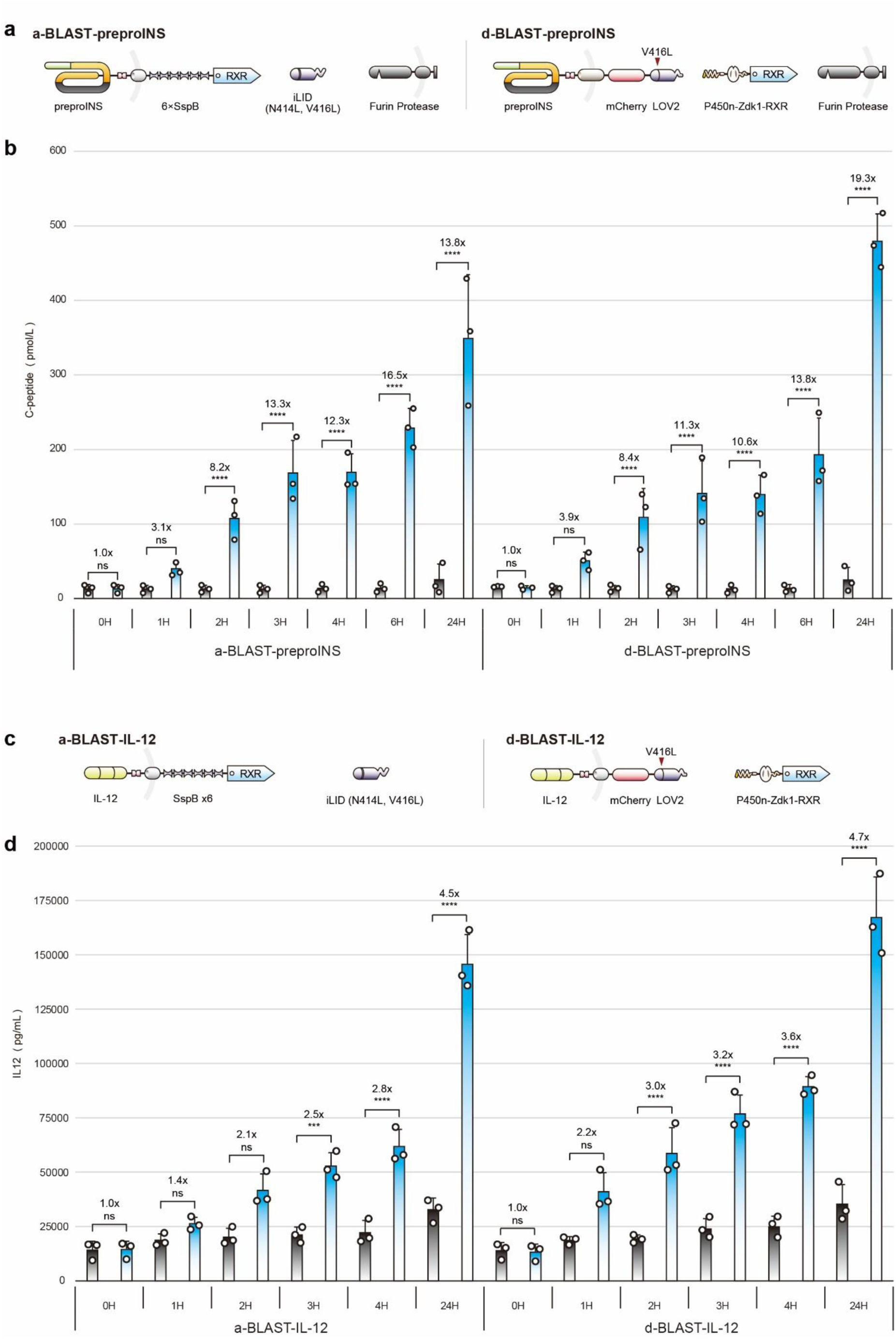
Secretion of therapeutic proteins with a-BLAST and d-BLAST. (a) The human preproinsulin (preproINS) construct contains a signal peptide, B-chain (yellow), C-peptide (dark gray), and A-chain (yellow), with engineered Furin cleavage sites flanking the C-peptide for maturation. To facilitate efficient processing within the Golgi apparatus, Furin protease was co-transfected with the BLAST modules. (b) Kinetic profiling of light-induced insulin secretion. Summary graphs of secreted C-peptide levels, quantified by ELISA, as a proxy for insulin secretion from a-BLAST (left) and d-BLAST (right). Both systems exhibited significant, time-dependent insulin release starting from 2 h of illumination (8.2-fold for a-BLAST, 8.4-fold for d-BLAST), reaching maximal induction at 24 h (13.8-fold for a-BLAST, 19.3-fold for d-BLAST). (c) Plasmid configurations for IL-12 secretion. Schematic of the heterodimeric cytokine IL-12-a-BLAST (left) and d-BLAST-IL-12 (right) constructs. (d) Kinetic profiling of light-induced IL-12 secretion. Summary graphs showing IL-12 secretion levels measured by ELISA. Significant secretion was observed starting at 3 h for a-BLAST (2.5-fold) and 2 h for d-BLAST (2.2-fold). At the 24 h time point, d-BLAST (4.7-fold) demonstrated a slightly higher dynamic range compared to a-BLAST (4.5-fold). Open circles represent individual measurements from three biologically independent samples. Data are presented as means ± S.D. Statistical significance was assessed using one-way ANOVA followed by Tukey’s multiple comparisons test (ns = not significant, \**P* < 0.05, \*\**P* < 0.01, \*\*\**P* < 0.001, \*\*\*\**P* < 0.0001).

Next, we extended the application to IL-12, a potent heterodimeric cytokine (comprising p35 and p40 subunits) critical for T-cell activation and cancer immunotherapy (Fig. 5c)^[38,39]^. Given its complex hetero dimeric structure, the successful secretion of IL-12 serves as a stringent test for the capacity of BLAST to handle multi-subunit proteins. Subsequent ELISA quantification revealed that d-BLAST triggered detectable IL-12 release within 2 h, whereas a-BLAST required 3 h to achieve comparable induction, further confirming the superior kinetic response of the dissociative system (Fig. 5d).

To provide an orthogonal validation of IL-12 release beyond ELISA-based quantification, and to specifically verify the biological activity of the optogenetically released cytokine, we employed IL-12 reporter cells (HEK-Blue™ IL-12), which express SEAP upon STAT4 activation (Supplementary Fig. 7). These assays demonstrated a robust 12.4- and 11.3-fold induction in IL-12-mediated signaling for a-BLAST and d-BLAST, respectively, compared to dark-state controls. Taken together, these findings collectively demonstrate that BLAST provides a versatile, on-demand platform for the controlled delivery of diverse therapeutic proteins, ranging from metabolic hormones to complex immunomodulators.

## 3 Discussion

The demand for precise spatiotemporal control over protein signaling has driven the development of various optogenetic post-translational toolkits in synthetic biology^[40–42]^. While traditional transcription-based circuits are effective for long-term modulation, their inherent 4-to 6 h kinetic lag limits their utility for mimicking rapid physiological events such as hormonal surges or neurotransmitter release^[7,43–45]^. Although recent advances in post-translational secretion, such as optoPASS^[23]^ and RELEASE^[20]^, have significantly improved temporal resolution, they largely rely on irreversible proteolytic cleavage to achieve secretion^[22,46,47]^.

Here, we present BLAST, a versatile optogenetic platform that achieves rapid and reversible control of protein secretion by harnessing light-tunable PPIs. The architectural hallmark of BLAST is its ability to override ER retention through steric masking rather than proteolytic cleavage. Crucially, this non-destructive mechanism endows BLAST with true reversibility—a feature fundamentally lacking in cleavage-based systems. As demonstrated by our alternating light/dark multi-toggling assays, while proteolytic switches lose control and continuously leak cargo in the dark following initial activation, BLAST achieves a highly regulated “staircase” secretion profile. The immediate re-establishment of the RXR masking upon light withdrawal completely and instantaneously arrests further cargo release. This capacity for repeated, strict ON/OFF switching provides the unprecedented temporal resolution required to emulate dynamic, pulsatile physiological signaling.

By situating the regulator module in the cytoplasm rather than the ER lumen, we bypassed the extensive protein engineering typically required to adapt optogenetic domains to the oxidative ER environment^[21,48,49]^. This modularity allows BLAST to be easily repurposed with alternative PPI pairs. Furthermore, our strategic selection of the RXR motif over the common KKMP motif provides a significant advantage. While KKMP-mediated retention is highly sensitive to its proximity to the membrane^[31,50,51]^, RXR motif allows for greater spatial flexibility^[26,31]^. In the d-BLAST system, this enabled us to incorporate mCherry as a structural spacer, without compromising retention efficiency, thereby facilitating the real-time visualization of cargo trafficking—a feature often challenging to implement in other systems.

A key advantage of the BLAST toolkit is the availability of two complementary operating modes, allowing users to tailor the system to specific experimental constraints. The associative a-BLAST, driven by iLID/SspB dimerization, offers exceptional stringency with minimal basal leakage in the dark state. This makes it the optimal choice for applications requiring a strict “OFF” switch, such as the regulation of potent cytokines or toxic payloads where background signaling must be minimized^[52]^. Conversely, the dissociative d-BLAST, based on LOV2/Zdk1, exhibits superior rapid-response kinetics, triggering significant secretion within 1 h of illumination. Although this sensitivity is accompanied by slightly higher basal secretion, d-BLAST is particularly advantageous for mimicking acute physiological bursts, such as insulin spikes or neurotransmitter release, where speed is paramount. Thus, BLAST offers a strategic trade-off: a-BLAST for high signal-to-noise precision, and d-BLAST for rapid temporal resolution.

In comparison to other light-triggered release systems, such as the UV-based PhoCl switch^[53,54]^, BLAST offers superior biocompatibility. While single-component UV switches are structurally compact, the high-energy UV radiation required for their activation is often cytotoxic and unsuitable for prolonged live-cell imaging^[55]^. BLAST utilizes 470 nm blue light, which is not only biocompatible but also permits high-resolution spatial patterning, as demonstrated by our slit-masking experiments. Unlike chemical induction systems like RUSH^[19]^, which are limited by the diffusion and slow washout of ligands, BLAST enables localized, on-demand protein delivery with high spatial precision.

Beyond its utility as a basic research tool for studying vesicular transport and protein trafficking, BLAST holds significant therapeutic potential. We successfully demonstrated the light-triggered release of functionally active insulin and IL-12 in HEK293T cells. Given the broad applicability of the BLAST platform across diverse mammalian cell lineages established earlier, this success suggest that BLAST can be used to engineer various non-specialized cells into programmable bio-factories for the localized delivery of therapeutic proteins. We anticipate that BLAST will serve as a foundational technique for synthetic biology, enabling the development of next-generation cell-based therapies with unprecedented spatiotemporal control.

## 4 Experimental Section

### 4.1 Design and construction of plasmid vectors

All plasmids used in this study are detailed in the Supplementary Information. Plasmids were constructed using standard molecular cloning techniques. DNA fragments were obtained from Addgene or synthesized (Bionics, Republic of Korea) and digested with restriction enzymes (New England Biolabs; NEB). Ligation was performed using T4 DNA Ligase (NEB, M0202S), and the resulting products were transformed into DH5α *Escherichia coli* competent cells (Enzynomics, Republic of Korea). PCR amplifications were performed using KOD Hot Start DNA Polymerase (Novagen) or Platinum SuperFi II PCR Master Mix (Invitrogen) according to the manufacturer’s protocols. All plasmid sequences were verified by Sanger sequencing (Bionics, Republic of Korea). Large-scale plasmid preparation was performed using 200 mL cultures. SnapGene software (Insightful Science) was used for vector design and sequence documentation.

### 4.2 Cell culture and plasmid DNA transfection

HEK293T, HeLa, HepG2, and BHK-21 cells were obtained from the American Type Culture Collection (ATCC). HEK-Blue™ IL-12 reporter cells were purchased from InvivoGen. All cell lines were confirmed to be mycoplasma-free and maintained in Dulbecco’s Modified Eagle’s Medium (DMEM; WELGENE, Republic of Korea) supplemented with 10% fetal bovine serum (FBS; Gibco, USA) and 1% penicillin-streptomycin (Gibco) at 37°C in a humidified 5% CO₂ atmosphere. For subculturing, cells were dissociated with 0.25% trypsin-EDTA, centrifuged at 123 × g for 1 m, and resuspended in fresh medium. Cells were seeded onto poly-D-lysine (Invitrogen)-coated 24-well plates at a density of 1 × 10^5^ cells per well. Cell viability and counting were assessed using a Countess automated cell counter (Invitrogen). HEK293T cells were transfected 24 h post-seeding using the jetOPTIMUS DNA transfection kit (Polyplus) following the manufacturer’s instructions (0.25 μg DNA and 0.25 μl reagent per well). HeLa, HepG2, and BHK-21 cells were transfected 24 h post-seeding using the PEI MAX DNA transfection kit (Polyscience) following the manufacturer’s instructions (0.5 μg DNA and 2 μg reagent per well).

### 4.3 Blue-light illumination

For optogenetic blue light irradiation of the cells as described in the figure, a 470-nm light-emitting diode (LED) panel (Green Energy Star) was installed inside a 37°C incubator maintained at 5% CO_2_. To prevent overheating via direct contact with the LEDs, sample culture plates were placed on three empty plates (total height, 6cm). The duty cycle of the LED panel was controlled by two serially connected electronic timers (IRT16-D, Han Seung). The light intensity was calibrated to 4.0 mW cm⁻² using a power meter (PM100D, Thorlabs). For dark conditions, plates were strictly shielded from ambient light using aluminum foil.

### 4.4 Chemical stimulation

Rapamycin (Tocris) was prepared as a 3 mM stock solution in DMSO (Sigma-Aldrich). For chemo-BLAST induction, cells were treated with the indicated concentration of rapamycin or an equivalent volume of DMSO as a vehicle control.

### 4.5 SEAP assay

For the quantification of SEAP, the cell culture supernatants were collected and centrifuged at 15,800 × *g* for 5 m. Endogenous alkaline phosphatase was heat-inactivated at 60°C for 1 h. A 40 μl aliquot of the supernatant was mixed with 100 μl of reaction buffer (5.1 M diethanolamine, L-homoarginine, and 10 mM MgCl₂) and incubated at 37°C for 10 m. Subsequently, 60 μl of pNPP substrate (Thermo Fisher Scientific) was added. Absorbance at 405 nm was monitored at 30 s intervals for 30 m using an Infinite F50 microplate reader (TECAN). Vmax values were calculated using Magellan software (TECAN).

### 4.6 Reversibility assay

To evaluate the dynamic, reversible control of protein secretion, HEK293T cells transfected with a-BLAST, d-BLAST, or NEW optoPASS constructs were seeded in 6-well plates. At 48 h post-transfection, the cells were subjected to a 9 h alternating light/dark cycle (3 h ON / 3 h OFF / 3 h ON). Blue light stimulation was delivered using a 470 nm LED panel with a pulsed duty cycle (2 s ON / 58 s OFF). During the dark phases, culture plates were strictly shielded from ambient light using aluminum foil. To track cumulative secretion profiles, 200 μL aliquots of culture supernatant were sampled from each well (initial volume: 2 mL) at designated time points (0, 3, 6, and 9 h). Immediately following each sampling, an equal volume (200 μL) of pre-warmed fresh medium was replenished to maintain a constant culture volume and minimize cellular stress. To precisely quantify the cumulative SEAP secretion and account for the serial dilution caused by the repeated sampling and replenishment process, the raw SEAP activity (*C*_*n*_, measured as Vmax) at each *n*-th time point was mathematically corrected. The corrected cumulative concentration (*C*_*corrected*,*n*_) was calculated using the following equation:

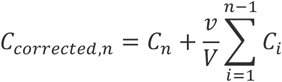

where *v* represents the sampled volume (200 μL), *V* represents the total culture volume in the well (2,000 μL), and the summation term accounts for the total mass of SEAP removed during all preceding sampling events.

### 4.7 Image processing and visualization

To visualize intercellular protein transfer, receiver cells (transfected with the reporter) and sender cells (transfected with BLAST) were prepared on separate coverslips and wells, respectively. The coverslip containing receiver cells was transferred to the sender cell well, placed with the cell-attached side facing downward to ensure proximity. Following 24 h of blue light stimulation (2 s ON / 58 s OFF), the coverslips were transferred to a new plate, fixed with 4% paraformaldehyde (CUREBIO) for 20 m, and washed with PBS. Samples were mounted using Crystal Mount (Biomeda). Confocal images were acquired using an LSM800 microscope (Zeiss) with a 40×/0.8 NA objective. For the slit-masking experiment (Supplementary Fig. 6), fluorescence scanning was performed using a Typhoon FLA 9500 (GE Healthcare).

### 4.8 Quantification of insulin and IL-12 secretion

For insulin quantification, HEK293T cells were co-transfected with the BLAST-preproinsulin constructs and Furin protease to enable maturation. Secreted human C-peptide levels in the supernatant were quantified using a Human C-peptide ELISA Kit (R&D Systems; Catalog #DICP00). Similarly, IL-12 secretion was measured using a Human IL-12 p70 ELISA Kit (R&D Systems; Catalog #D1200) according to the manufacturer’s instructions. To utilize HEK-Blue™ IL-12 reporter cells for measuring optogenetically induced IL-12 secretion, a 5-day experimental protocol was employed (Supplementary Fig. 7). HEK293T cells and HEK-Blue™ IL-12 reporter cells were seeded into 24-well plates on Day 1 and Day 3, respectively. Twenty-four h after seeding, HEK293T cells were transfected with BLAST-IL-12 (Day 2). Twenty-four h post-transfection, cells were cultured for 24 h under dark or blue light conditions (Day 3). On Day 4, culture supernatants containing secreted IL-12 were collected and transferred to HEK-Blue™ IL-12 reporter cells seeded on Day 3. After 24 h of incubation to promote signal transduction, SEAP activity was measured on Day 5.

### 4.9 Statistical analysis

Data are presented as mean values ± S.D. from at least three biologically independent experiments. Statistical comparisons were performed using one-way or two-way ANOVA followed by Tukey’s multiple comparisons test using GraphPad Prism or Microsoft Excel. Adjusted P-values are denoted as: ns (not significant), *p < 0.05, **p< 0.0 1, ***p < 0.001, and ****p < 0.0001.

### 4.10 Manuscript Preparation and AI Usage

During the preparation of this manuscript, the authors utilized Gemini (Google) for English language editing, stylistic polishing, and formatting of the text. After using this tool, the authors carefully reviewed and extensively edited the content. The authors take full responsibility for the final content, underlying data, and scientific accuracy of this publication.

## Acknowledgements

This study was supported by grants from the National Research Foundation of Korea (grant nos. RS-2020-NR051270 and RS-2024-00340694 to D.L.). We thank all members of the laboratory for their critical discussions and comments.

## Author Contributions

M.S., S.L., and D.L. developed the concept. M.S., S.L., and M.C. designed and implemented the methodology and performed the experiments. M.S., S.L., M.C., and D.L. performed data curation and visualization. D.L. supervised the project and secured funding. M.S., S.L., and D.L. wrote the manuscript. All authors reviewed and edited the manuscript.

## Data Availability Statement

The data that support the findings of this study are available from the corresponding author upon reasonable request.

---

**Supplementary figure 1.**
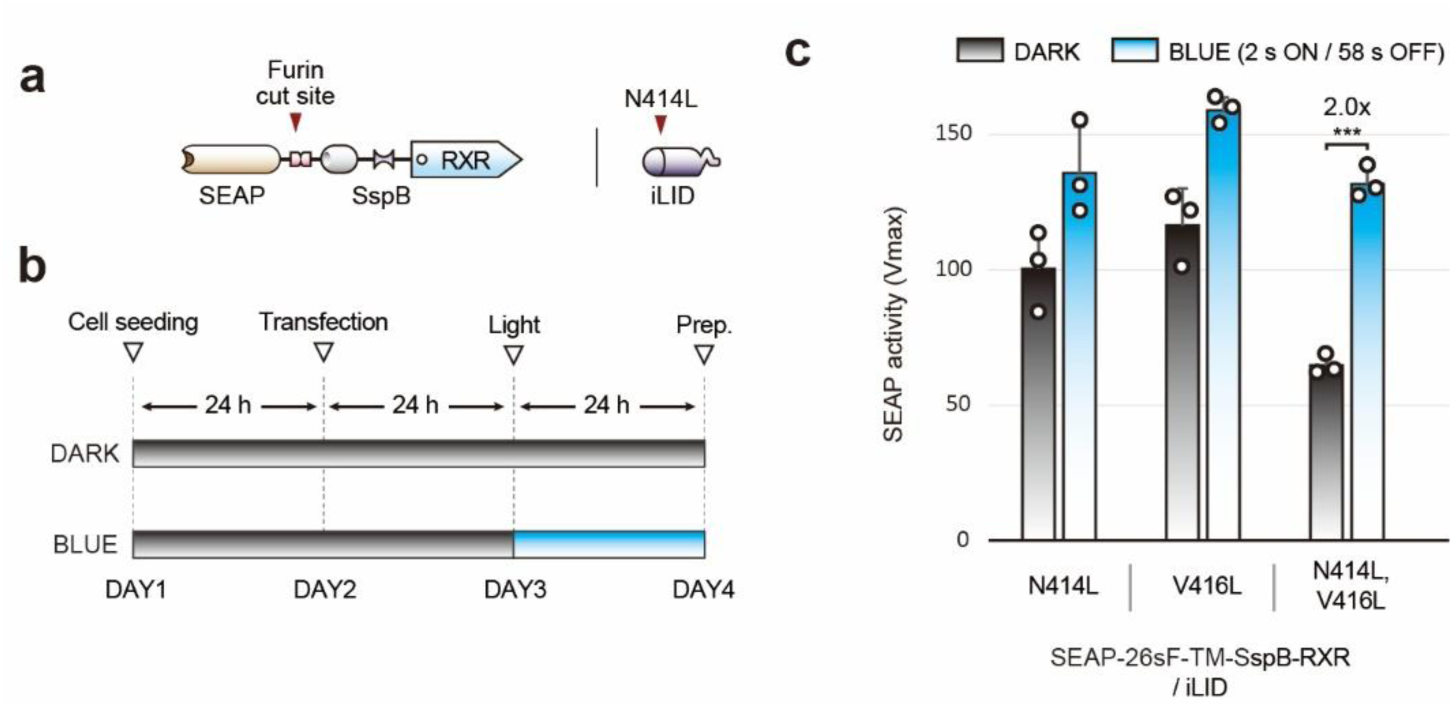
Optimization of the a-BLAST prototype using iLID mutants. (a) Schematic of the prototype a-BLAST components. The cargo module (upper) consists of SEAP, a Furin cleavage site, a transmembrane (TM) domain, a single SspB domain, and an RXR retention motif. The regulator module (lower) expresses iLID variants (e.g., N414L) derived from iLID, engineered to modulate dark-state recovery kinetics. (b) Experimental timeline for SEAP secretion assay. Schematic of the 4-day protocol. Cells were seeded (Day 1), transfected (Day 2), and incubated for 24 h. Blue light stimulation (Day 3) was applied for 24 h using a pulsed duty cycle (2 s ON / 58 s OFF) before supernatant analysis (Day 4). (c) Screening of iLID affinity mutants. Comparison of light-induced SEAP secretion using different iLID variants (N414L, V416L, and N414L/V416L) paired with the prototype cargo (1× SspB). While single mutants exhibited high basal leakage, the double mutant (N414L, V416L) showed a statistically significant fold change (2.0-fold), identifying it as the optimal candidate for further engineering. Open circles represent individual measurements from three biologically independent samples. Data are presented as means ± S.D. Statistical significance was assessed using one-way ANOVA followed by Tukey’s multiple comparisons test (ns = not significant, \**P* < 0.05, \*\**P* < 0.01, \*\*\**P* < 0.001, \*\*\*\**P* < 0.0001).

**Supplementary Figure 2.**
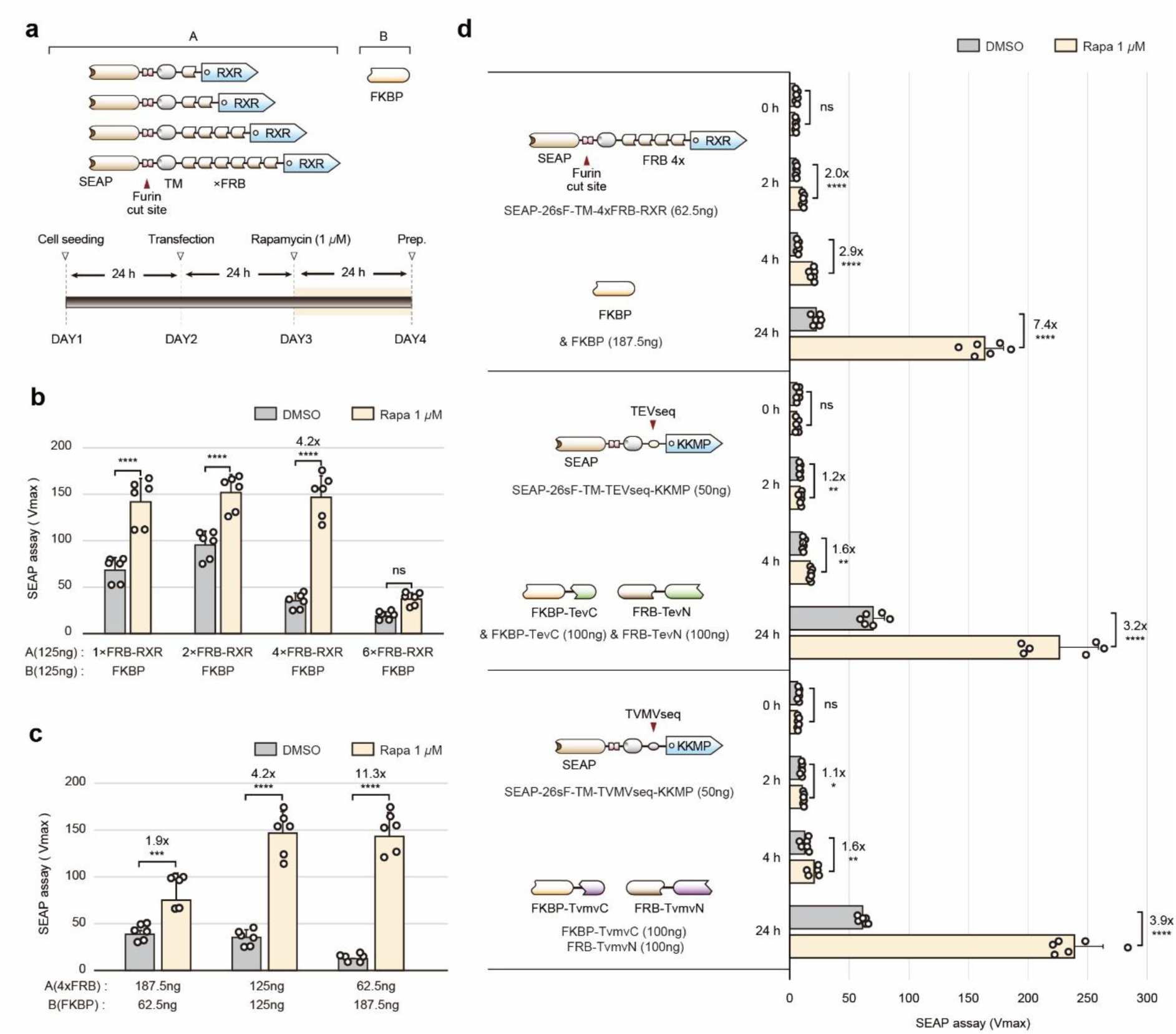
Benchmarking of chemogenetic BLAST against the RELEASE system. (a) Design of the chemogenetic BLAST (chemo-BLAST) and experimental timeline. Schematic of the plasmid configurations (upper) and the 4-day experimental schedule (lower). To enable a direct mechanistic comparison independent of light stimulation, the optogenetic modules (iLID/SspB) were replaced with the chemically inducible rapamycin-binding domains (FKBP/FRB). The cargo (A) contains varying repeats of FRB (1×, 2×, 4×, and 6×), and the regulator (B) consists of FKBP. Rapamycin (1 μM) was used as the inducer. (b) Optimization of FRB valency. Summary graph of SEAP secretion levels with varying FRB repeats. The construct containing 4× FRB repeats exhibited the most effective switching performance (4.2-fold induction). (c) Optimization of transfection ratios. Using the 4× FRB construct, SEAP levels were measured across different cargo (A) to regulator (B) ratios. The 1:3 ratio (62.5 ng : 187.5 ng) yielded the highest dynamic range (11.3-fold). (d) Head-to-head kinetic comparison with the RELEASE system. The optimized chemo-BLAST was compared against the RELEASE system, which employs split proteases (TEVp or TVMVp) to cleave a KKMP ER retention motif. Chemo-BLAST induced significant protein secretion starting at 2 h and achieved a higher fold change (7.4-fold at 24 h) compared to the RELEASE systems (3.2-fold for TEVp, 3.9-fold for TVMVp), demonstrating superior kinetics and signal-to-noise ratio. Open circles represent individual measurements from three biologically independent samples. Data are presented as means ± S.D. Statistical significance was assessed using one-way ANOVA followed by Tukey’s multiple comparisons test (ns = not significant, \**P* < 0.05, \*\**P* < 0.01, \*\*\**P* < 0.001, \*\*\*\**P* < 0.0001).

**Supplementary Figure 3.**
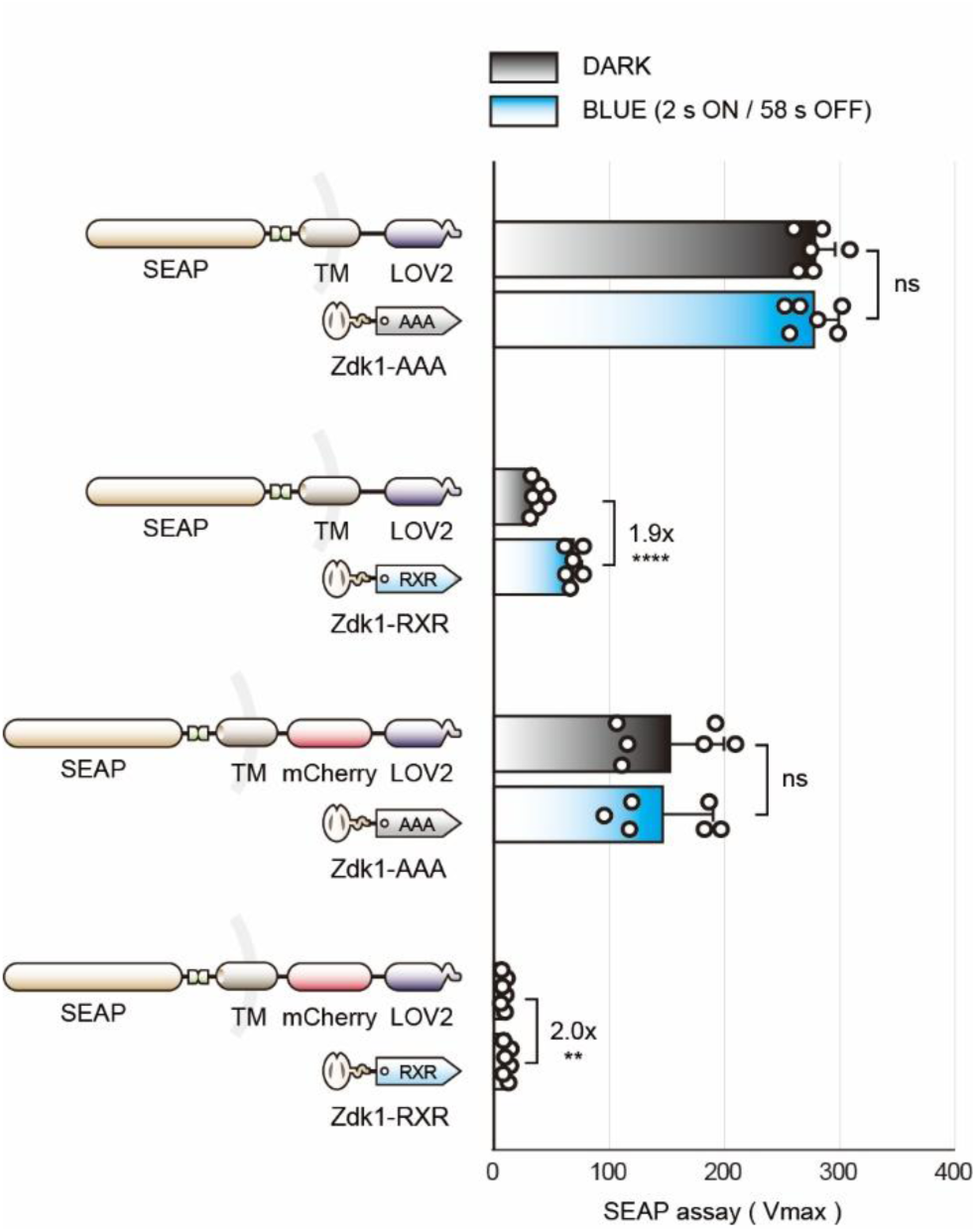
Impact of the mCherry spacer on d-BLAST basal retention. To evaluate the structural necessity of the mCherry spacer, HEK293T cells were co-transfected with the ER-anchored regulator (P450n-Zdk1-RXR or the non-functional Zdk1-AAA control) and d-BLAST cargo variants either lacking (SEAP-Furin-TM-LOV2) or containing (SEAP-Furin-TM-mCherry-LOV2) the mCherry spacer. The Zdk1-AAA mutant, lacking the retention motif, served as a negative control for constitutive secretion. Notably, the construct lacking mCherry exhibited higher basal secretion in the dark compared to the mCherry-containing variant. This suggests that the mCherry spacer improves retention efficiency by optimizing the spatial distance of the LOV2 binding domain from the ER membrane. Open circles represent individual measurements from three biologically independent samples. Data are presented as means ± S.D. Statistical significance was assessed using one-way ANOVA followed by Tukey’s multiple comparisons test (ns = not significant, \**P* < 0.05, \*\**P* < 0.01, \*\*\**P* < 0.001, \*\*\*\**P* < 0.0001).

**Supplementary Figure 4.**
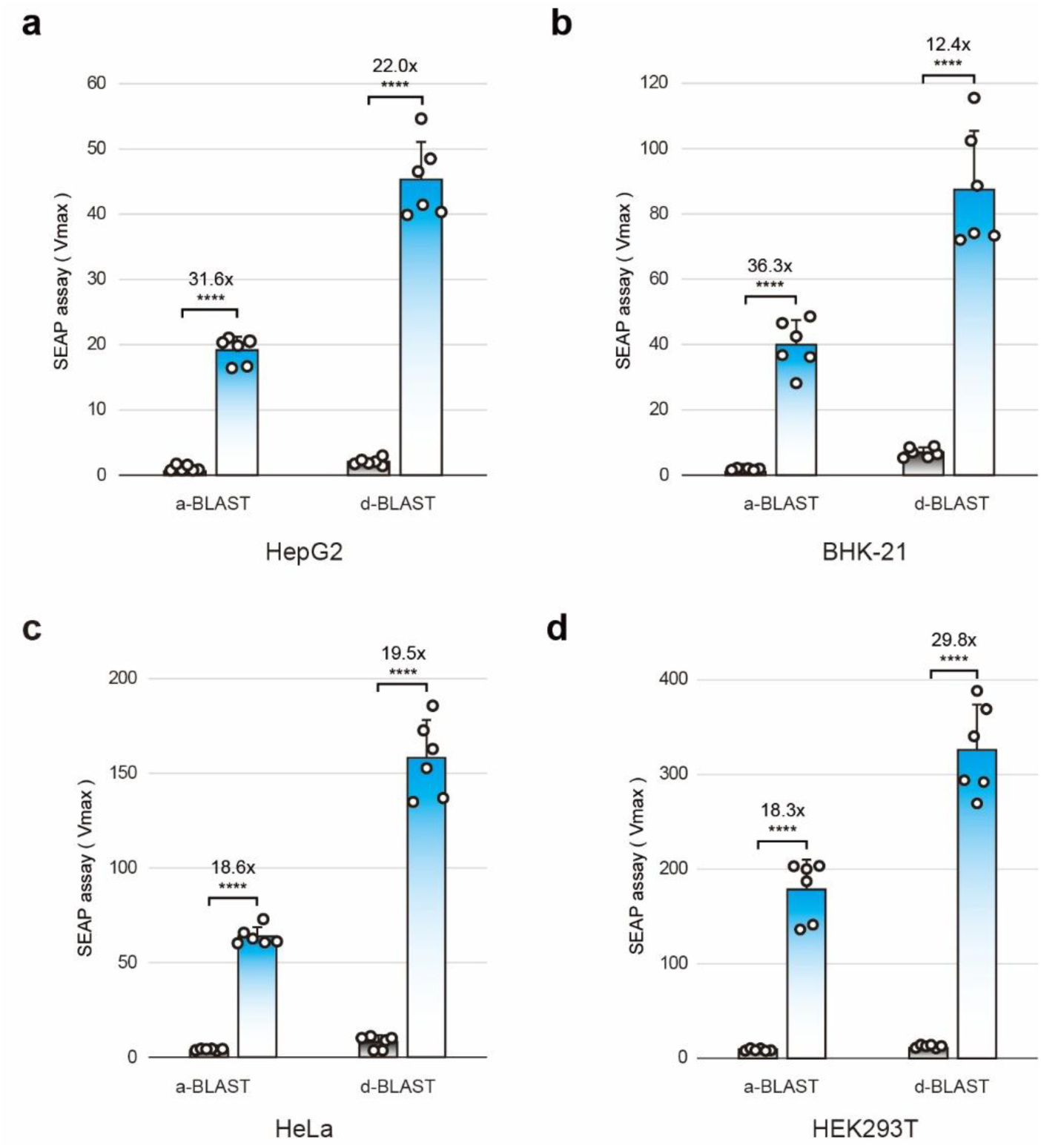
Validation of BLAST functionality across diverse mammalian cell lineages. Quantification of SEAP secretion levels in the supernatants of various mammalian cell lines following transient transfection with a-BLAST or d-BLAST. The assay was performed in (a) HepG2 (human liver cancer), (b) BHK-21 (baby hamster kidney), (c) HeLa (human cervical cancer), and (d) HEK293T (human embryonic kidney) cells. Both a-BLAST and d-BLAST exhibited robust, light-dependent protein secretion with high dynamic ranges across all tested cell types, confirming the broad applicability of the toolkit. Open circles represent individual measurements from three biologically independent samples. Data are presented as means ± S.D. Statistical significance between dark and blue light conditions was assessed using one-way ANOVA followed by Tukey’s multiple comparisons test (\*\*\*\**P* < 0.0001).

**Supplementary Figure 5.**
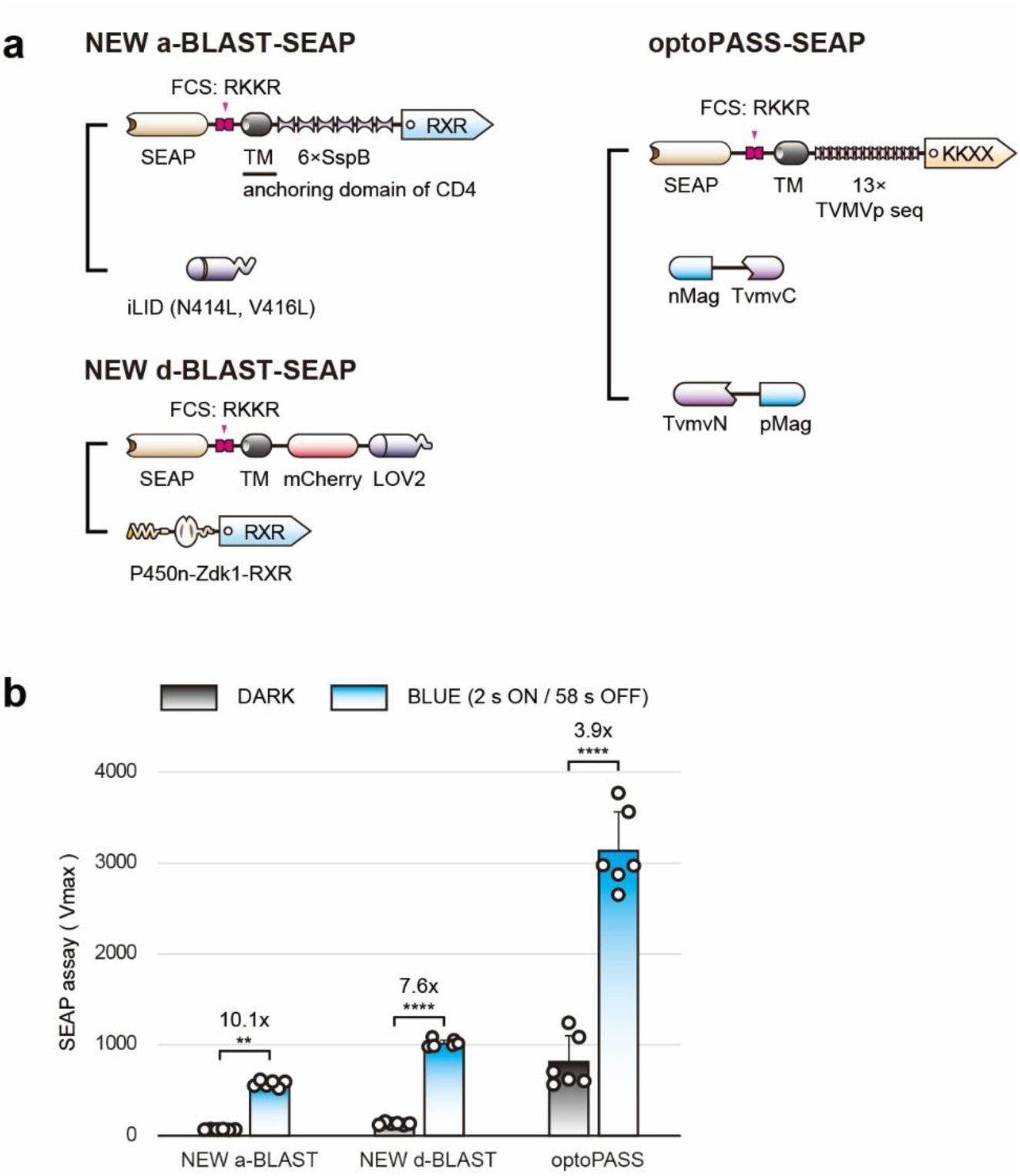
Benchmarking of NEW BLAST variants containing optoPASS-derived domains. (a) Design of chimeric NEW BLAST constructs. Schematic representation of the engineered NEW a-BLAST-SEAP and NEW d-BLAST-SEAP systems alongside SEAP-optoPASS. To strictly control domain-specific effects, the native Furin cleavage site (FCS) and transmembrane (TM) domain of the original BLAST systems were replaced with the influenza virus-derived sequence (RRRKKR/GL) and the CD4 anchoring domain, respectively, matching the exact configuration used in optoPASS. (b) Summary graph of SEAP secretion levels. While optoPASS showed higher absolute secretion levels, it suffered from high basal leakage in the dark. Consequently, the NEW a-BLAST (10.1-fold) and NEW d-BLAST (7.6-fold) variants exhibited superior dynamic ranges compared to optoPASS (3.9-fold). This confirms that the superior signal-to-noise ratio of BLAST is intrinsic to its reversible retention mechanism, rather than dependent on specific structural domains. Open circles represent individual measurements from three biologically independent samples. Data are presented as means ± S.D. Statistical significance between dark and blue light conditions was assessed using one-way ANOVA followed by Tukey’s multiple comparisons test (ns = not significant, \**P* < 0.05, \*\**P* < 0.01, \*\*\**P* < 0.001, \*\*\*\**P* < 0.0001).

**Supplementary Figure 6.**
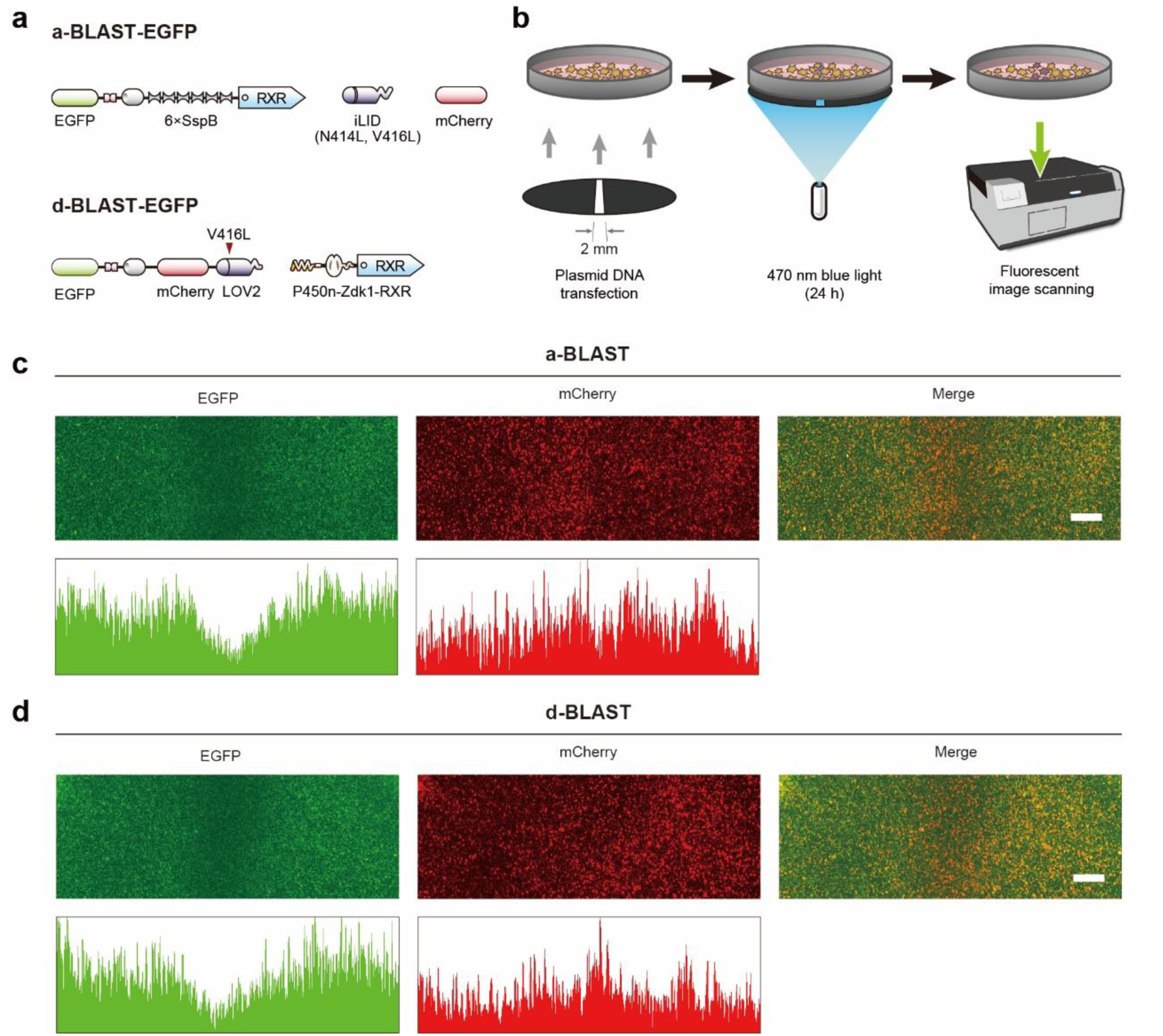
High-resolution spatial control of protein secretion via slit-masking. (a) Schematic of EGFP-fused BLAST constructs. Diagrams of the a-BLAST-EGFP and d-BLAST-EGFP vectors used for spatial patterning experiments. EGFP serves as the fluorescent cargo to visualize intracellular depletion upon secretion. (b) Experimental workflow for slit-guided photopatterning. Schematic illustrating the procedure: Transfected cell monolayers were covered with a black mask containing a 2 mm slit and exposed to blue light (470 nm) for 24 h. (c) and (d) Visualization of spatially restricted secretion. Representative fluorescence images (upper) and corresponding intensity profile plots (lower) for a-BLAST (c) and d-BLAST (d). Left (EGFP): Shows the intracellular cargo. A distinct dark band (fluorescence depletion) is visible in the slit region, indicating light-triggered secretion. Middle (mCherry): Shows the co-transfected mCherry cell marker, which serves as a non-secreted reference. The signal remains constant across the slit, confirming uniform cell density. Right (Merge): Merged image showing the spatial contrast between the secreted cargo (green) and the retained marker (red). Scale bar: 2 mm.

**Supplementary Figure 7.**
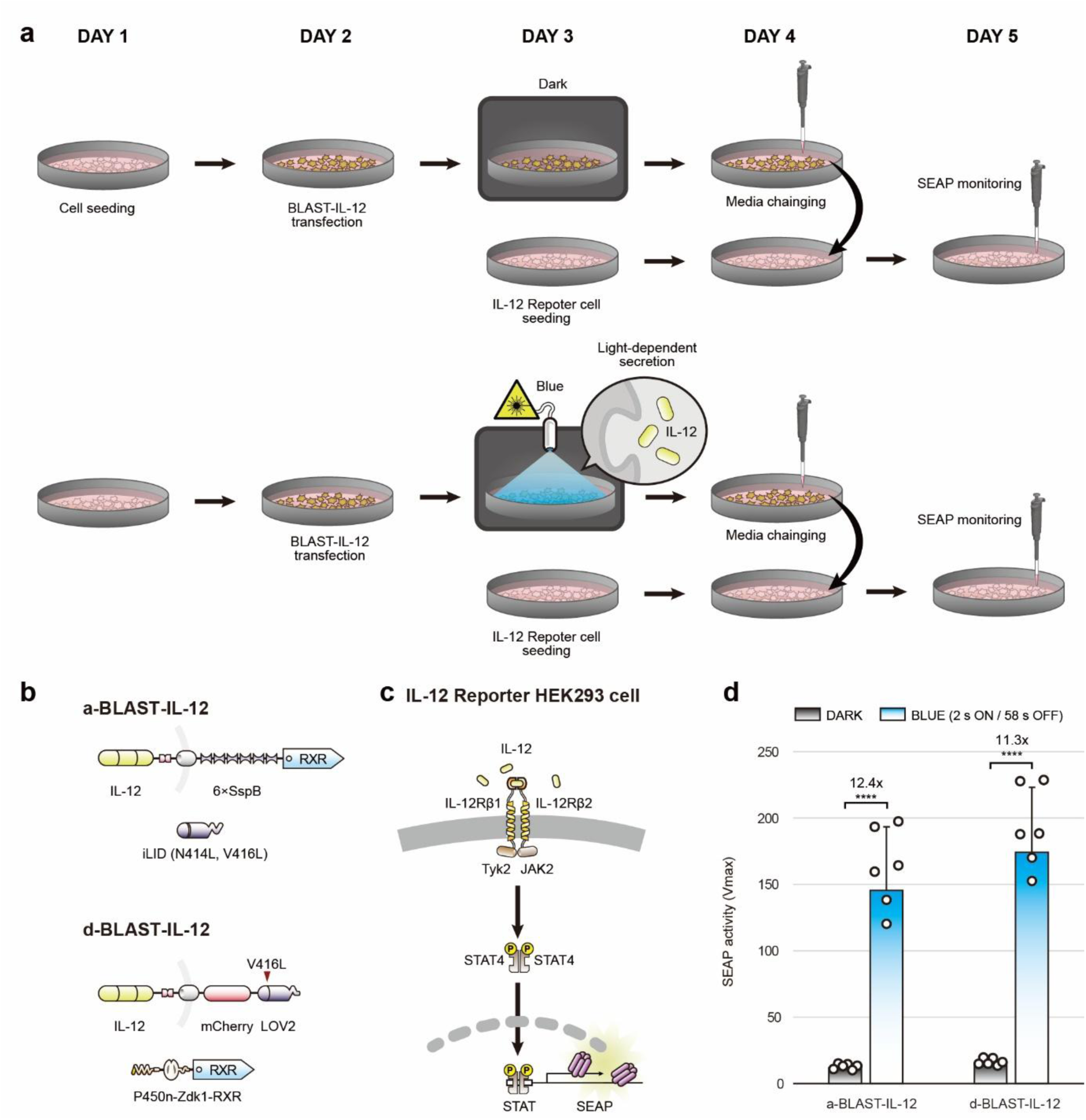
Functional validation of optogenetically secreted IL-12 using a reporter cell assay. (a) Schematic of the experimental workflow for bioactivity validation. The protocol spans 5 days. HEK293T cells transfected with BLAST-IL-12 were subjected to dark or blue light conditions for 24 h (Day 3). On Day 4, the conditioned media containing secreted IL-12 was harvested and transferred to HEK-Blue™ IL-12 reporter cells (seeded on Day 3). After a 24 h incubation to allow for signal transduction, SEAP activity was quantified (Day 5). (b) Plasmid configurations. Schematics of the a-BLAST-IL-12 (upper) and d-BLAST-IL-12 (lower) constructs used for the assay. (c) Illustration of the JAK-STAT signaling pathway in the reporter cells. Binding of secreted IL-12 to the IL-12 receptor complex activates Tyk2/JAK2, leading to STAT4 phosphorylation. Phosphorylated STAT4 dimerizes and translocates to the nucleus to induce SEAP expression. (d) Summary graph of SEAP activity induced by the conditioned media. The results confirm that both systems secrete biologically active IL-12 upon blue light stimulation, exhibiting robust fold changes (12.4-fold for a-BLAST and 11.3-fold for d-BLAST) compared to the dark control. Data are presented as means ± S.D. Statistical significance was assessed using one-way ANOVA followed by Tukey’s multiple comparisons test (\*\*\*\**P* < 0.0001).

